# Sex differences in gene expression in the human fetal brain

**DOI:** 10.1101/483636

**Authors:** Heath E. O’Brien, Eilis Hannon, Aaron R. Jeffries, William Davies, Matthew J. Hill, Richard J. Anney, Michael C. O’Donovan, Jonathan Mill, Nicholas J. Bray

**Affiliations:** MRC Centre for Neuropsychiatric Genetics & Genomics, Division of Psychological Medicine & Clinical Neurosciences, Cardiff University School of Medicine, United Kingdom; University of Exeter Medical School, United Kingdom; School of Psychology, Cardiff University, United Kingdom

**Keywords:** Sex biases, brain, gene expression, developmental, prenatal, autism

## Abstract

Widespread structural, chemical and molecular differences have been reported between the male and female human brain. Although several neurodevelopmental disorders are more commonly diagnosed in males, little is known regarding sex differences in early human brain development. Here, we used RNA sequencing data from a large collection of human brain samples from the second trimester of gestation (N = 120) to assess sex biases in gene expression within the human fetal brain. In addition to 43 genes (102 Ensembl transcripts) transcribed from the Y-chromosome in males, we detected sex differences in the expression of 2558 autosomal genes (2723 Ensembl transcripts) and 155 genes on the X-chromosome (207 Ensembl transcripts) at a false discovery rate (FDR) < 0.1. Genes exhibiting sex-biased expression in human fetal brain are enriched for high-confidence risk genes for autism and other developmental disorders. Male-biased genes are enriched for expression in neural progenitor cells, whereas female-biased genes are enriched for expression in Cajal-Retzius cells and glia. All gene- and transcript-level data are provided as an online resource (available at http://fgen.psycm.cf.ac.uk/FBSeq1) through which researchers can search, download and visualize data pertaining to sex biases in gene expression during early human brain development.

## INTRODUCTION

Sex differences have been reported in the regional volumes (Ruigrok et al. 2014; Ritchie et al. 2018) connectivity (Ingalhalikar et al. 2014) and chemistry (Nishizawa et al. 1997; Laakso et al. 2002) of the adolescent and adult human brain. Sex biases are also observed in the prevalence and presentation of many human disorders of the central nervous system (Zagni et al. 2016). These include conditions with early neurodevelopmental origins such as autism spectrum disorders (ASD) and intellectual disability (Werling and Geschwind 2013; Polyak et al. 2015), the former diagnosed at a four-fold excess in males. Studies in animals have highlighted the importance of the pre-natal period in sexual differentiation of the brain and in establishing later sex-biased behaviours (Phoenix et al. 1959; Arnold 2009). However, little is currently known regarding sex differences in human fetal brain development.

Transcriptomic studies, executed through microarray and more recently RNA sequencing (RNA-Seq) technology, provide a powerful means of assessing the molecular basis of sex differences. Such studies have revealed sex biases in autosomal gene expression as well as in the expression of genes on the sex chromosomes in the postnatal human brain (Weickert et al. 2009; Kang et al. 2011; Trabzuni et al. 2013; Xu et al. 2014; Mayne et al, 2016; Shi et al. 2016; Werling et al. 2016). Although sex biases in gene expression have also been reported in the prenatal human brain (Reinius and Jazin 2009; Kang et al. 2011; Shi et al. 2016; Werling et al. 2016), studies exploring this to date have been limited by small sample sizes (total N ≤ 20) and widely varying gestational ages, with all but one (Shi et al. 2016) relying on microarray-based measures of gene expression.

In the present study, we examined sex differences in gene expression using RNA sequencing data from a large (N = 120) collection of brain samples from the second trimester of gestation (O’Brien et al. 2018). Our combination of greatly increased sample size and deep RNA sequencing allowed us to identify thousands of previously undetected sex biases in gene expression in the human prenatal brain, at the individual transcript as well as whole gene level. We provide our data as a searchable online resource, available at http://fgen.psycm.cf.ac.uk/FBSeq1.

## RESULTS

### Sex biases in gene expression in the human fetal brain

We analysed RNA sequencing data from 120 human fetal brains (12-19 post-conception weeks [PCW]), generated through a previous study (O’Brien et al. 2018). Abundances of individual Ensembl transcripts were quantified using kallisto (Bray et al. 2016), with measures of each gene based on summation of counts assigned to all annotated transcripts for that gene. We thus derived expression measures for 94,969 expressed Ensembl transcripts, annotated to 31,378 genes. We compared individual gene- and transcript-level expression between males (N = 70; mean age = 14.3 PCW; age range = 12 – 19 PCW) and females (N = 50; mean age = 14.2 PCW; age range = 12 – 19 PCW), controlling for known variables as well as hidden confounders (see Methods), using DESeq2 (Love et al. 2014).

Analyses at the gene level indicated 2756 Ensembl genes that were differentially expressed between male and female prenatal brain at a false discovery rate [FDR] < 0.1. Of these, 1468 genes exhibited higher expression in males and 1288 genes were more highly expressed in females (Supplementary Table S1). Genes with higher expression in males included 1377 located on autosomes and 48 on the X-chromosome, as well as 43 genes expressed from the Y-chromosome. Genes with higher expression in females included 1181 located on autosomes and 107 genes mapping to the X-chromosome. Only 20% (507) of the identified autosomal genes, 36% (56) of the X-chromosome genes and 56% (24) of the Y-chromosome genes have previously been reported to show sex-biased expression in the fetal or adult human brain (Supplementary Table S2).

Analyses at the individual transcript level identified 3032 Ensembl transcripts with sex-biased expression (FDR < 0.1), of which 1389 were more highly expressed in males and 1643 more highly expressed in females (Supplementary Table S3). Transcripts with higher expression in males included 1243 located on autosomes and 44 derived from the X-chromosome, as well as 102 transcripts expressed from the Y-chromosome. Transcripts with higher expression in females included 1480 located on autosomes and 163 on the X-chromosome. We detected sex biases in 1220 autosomal transcripts annotated to 1095 genes that did not exhibit pronounced sex differences (FDR < 0.1) at the gene level. These transcript-specific autosomal sex differences were found to particularly manifest when the affected transcript(s) were not the predominant RNA species of the gene (e.g. Figure 1).

**Figure 1.**
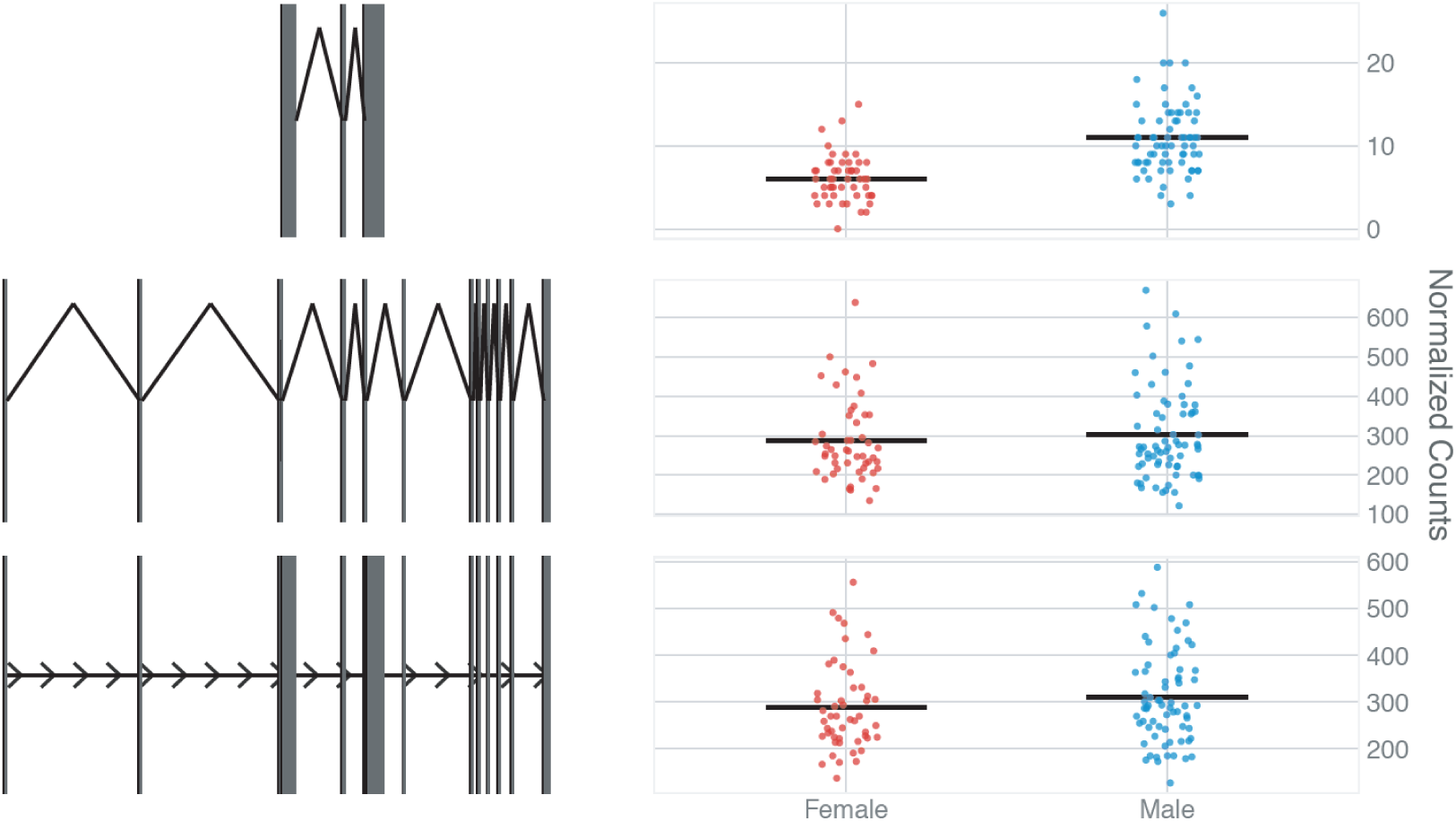
Transcript-specific sex-bias in expression of the autosomal gene *AKR1B15*. Significant male-bias is observed in the expression of a less abundant *AKR1B15* transcript ENST00000457545 (FDR = 4.8 × 10^−12^; top), but not in a more abundant *AKR1B15* transcript (ENST00000467545; FDR = 0.68; middle) or in *AKR1B15* expression at the summated gene level (FDR = 0.17; bottom).

### Sex-biased gene expression from the X-chromosome

X-chromosome inactivation (XCI) is the process by which transcription from one of the two X-chromosomes in female cells is silenced to balance gene expression dosage with XY males. Recent data indicate that at least 23% of X-chromosome genes escape XCI to some degree (Tukiainen et al. 2017). Twenty-six of the 107 X-chromosome genes that exhibited female-biased expression in fetal brain are reported by Tukiainen and colleagues to escape XCI, and a further 11 to show variable escape status (Figure 2, Supplementary Table S4). We note significant female biases (FDR < 0.1) in the expression of an additional 43 genes for which XCI status is either classed as unknown or is not reported in the Tukiainen et al. study (Figure 2; Supplementary Table S4), which could include novel candidates for XCI escape in the developing human brain. Eleven of the 48 X-chromosome genes exhibiting higher expression in males (FDR < 0.1) map to the PAR1 pseudoautosomal region, which showed a general male-bias in expression (Figure 2; Supplementary Table S4). Although the pseudoautosomal regions of the X-chromosome are believed to also escape XCI, our findings are consistent with data from adult human tissues (Tukiainen et al. 2017), indicating only partial dosage compensation in females.

**Figure 2.**
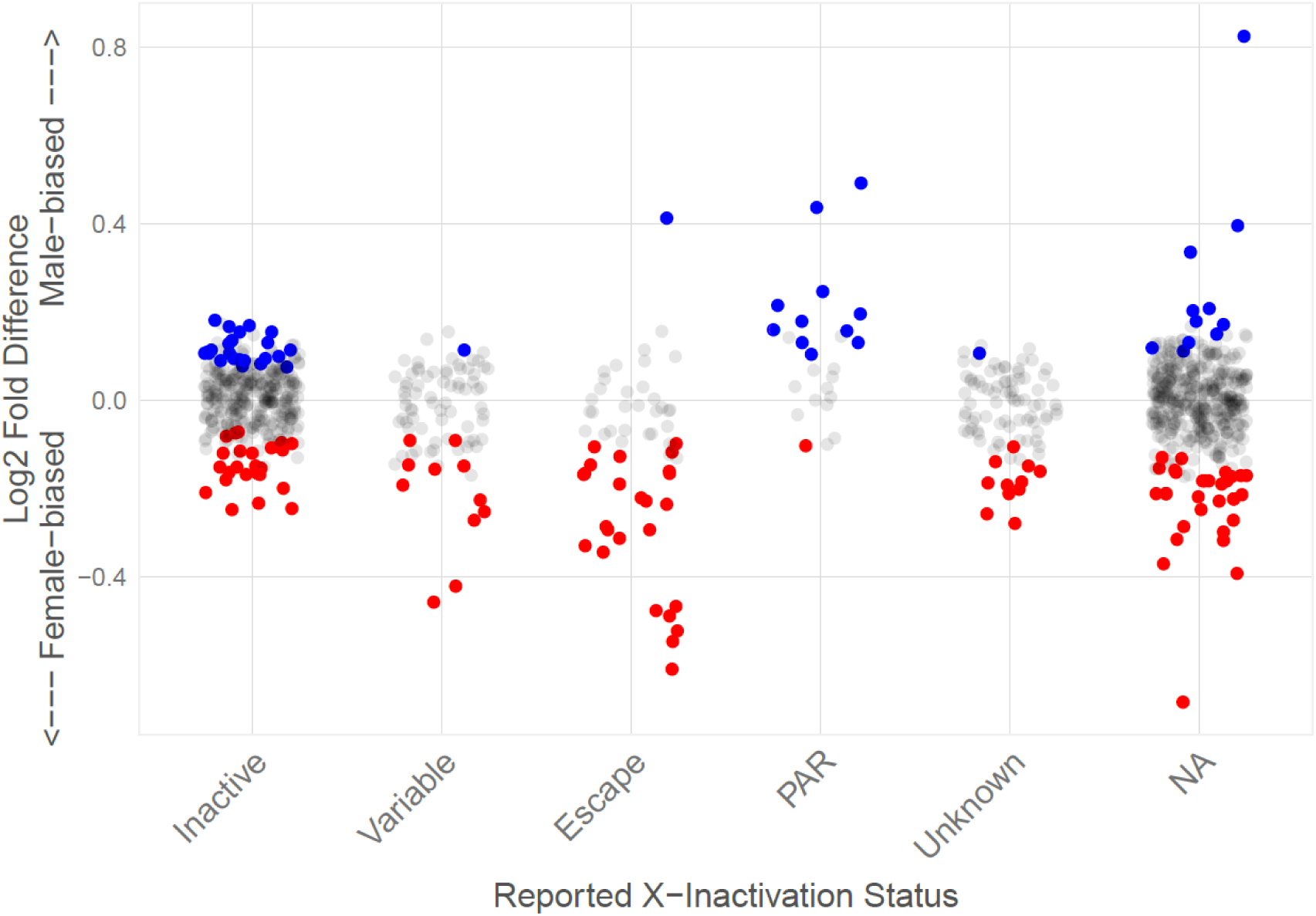
Sex biases in X-chromosome gene expression in relation to reported X-chromosome inactivation status (Tukiainen et al. 2017). Red data points indicate female-biased genes, while blue data points indicate male-biased genes (FDR < 0.1). Gray data points indicate X-chromosome genes that do not exhibit sex-biased expression at an FDR < 0.1. PAR = pseudoautosomal regions; NA = not reported in the study of Tukiainen and colleagues.

### Sex-biased expression of genes implicated in neurodevelopmental disorders

We detected expression of all 79 X-chromosome genes listed by Piton and colleagues (2013) as confirmed X-linked intellectual disability (XLID) loci, consistent with their involvement in early human brain development. In addition to the increased vulnerability to X-chromosome gene disruption in XY males, we note that 9 of these well-supported XLID genes (*KDM5C*, *NLGN4X*, *OFD1, SYN1, PTCHD1, IL1RAPL1, TSPAN7, CASK* and *RAB39B*) displayed higher baseline expression in the female prenatal brain (FDR < 0.1), while 8 (*FTSJ1*, *DLG3*, *ARHGEF9*, *FMR1*, *DKC1*, *RBM10*, *BCOR*, *FLNA*) exhibited higher expression in males. Three of the 10 most significantly female-biased genes (*STS*, *PUDP* and *NLGN4X*) map within a small region on chromosome Xp22.3 where deletions are associated with a substantially increased risk of developmental disorder and associated traits in males (Chatterjee et al. 2016). We detected expression of all but 2 (*MFRP* and *OR52M1*) of the 65 risk genes for autism (FDR ≤ 0.1) identified by Sanders and colleagues (2015). These high-confidence autism risk genes were significantly enriched for sex-biases in expression in the human fetal brain (18% sex-biased, compared with 9% of expressed genes overall, *P* = 0.004), with 7 autosomal genes (*ACHE*, *CHD2*, *KDM6B*, *KMT2C*, *PHF2*, *POGZ*, *SYNGAP1*) displaying higher expression in males and 5 autosomal genes (*ADNP*, *KAT2B*, *RANBP17*, *SCN2A*, *USP45*) exhibiting higher expression in females (at FDR < 0.1). Similarly, we found that 75 out of 76 autosomal genes exhibiting a genome-wide significant excess of damaging *de novo* mutations in developmental disorders (Deciphering Developmental Disorders Study, 2017) are expressed in fetal brain. These too were enriched for sex biases in gene expression, with 13 genes (*ADNP*, *MED13L*, *TCF4*, *EP300*, *FOXP1*, *CDK13*, *TBL1XR1*, *KAT6B*, *CHD2*, *POGZ*, *EHMT1*, *CTCF*, *AUTS2*) displaying higher expression in males and 3 genes (*SCN2A*, *COL4A3BP*, *DNM1*) exhibiting higher expression in females (21% sex-biased, *P* = 0.0004).

### Enrichment of sex-biased autosomal genes in functional categories and fetal cell types

Sex differences in autosomal gene expression could reflect cellular differences between the sexes that are a distal consequence of signalling through the sex chromosomes. As an initial exploration of the biological significance of our findings, we tested whether male- and female-biased autosomal genes (FDR < 0.1) were enriched for annotation to specific Gene Ontology (GO) terms. Autosomal genes that exhibited higher expression in males were enriched for GO terms including ‘chromosome organization’ (fold-enrichment = 2.5; Bonferroni-corrected *P* = 9.8 × 10^−23^) and ‘cell cycle’ (fold-enrichment = 1.9; Bonferroni-corrected *P* = 2.9 × 10^−11^), whereas those that were more highly expressed in females were notably enriched for synaptic processes, with ‘synaptic signaling’ the most significant term (fold-enrichment = 2.8; Bonferroni-corrected *P* = 1.2 × 10^−12^) (Supplementary Tables S5 and S6). These enrichments could reflect differences in cellular composition between the male and female brain at this developmental time-point. To explore this hypothesis, we tested for enrichment of sex-biased genes (FDR < 0.1) within gene sets that have recently been reported to distinguish cell types in the human fetal brain through single cell RNA sequencing (Fan et al. 2018; Zhong et al. 2018). Consistent with our finding that male-biased genes are enriched for involvement in the cell cycle, male-biased genes are also enriched among those that are reported to show higher expression in neural progenitor cells (Bonferroni-corrected *P* = 0.0038). In contrast, female-biased genes are notably enriched among those distinguishing Cajal-Retzius cells (Bonferroni-corrected *P* = 3.2 × 10^−49^), as well as for genes more highly expressed in oligodendrocyte precursor cells (Bonferroni-corrected *P* = 2.7 × 10^−6^), astrocytes (Bonferroni-corrected *P* = 0.025) and glia in general (Bonferroni-corrected *P* = 0.005).

## DISCUSSION

We present the largest analysis of transcriptional sex differences in the human prenatal brain to date, identifying sex-biases in the expression of approximately 9% of transcribed genes.

Our data illuminate molecular differences between males and females that are likely to govern as well as reflect the early sexual differentiation of the human brain. The former might include expression of 43 genes from the Y-chromosome in the male fetal brain, as well as sex-biased expression of X-chromosome genes in females. Sex biases in autosomal gene expression could reflect hormonal influences, the direct action of transcription factors transcribed from the sex chromosomes or cellular differences between males and females that are secondary to these factors. For example, studies in rodents indicate that fetal testosterone, through its aromatization to estradiol, could plausibly account for the increased cellular proliferation in the male prenatal brain suggested by our data (Martínez-Cerdeño et al. 2006; Bowers et al. 2010).

Our detection in the fetal brain of numerous genes implicated in XLID, ASD and other developmental disorders is consistent with a prenatal component to these conditions. In addition, we find that many of these genes display sex-biased expression in the human fetal brain, which could modulate the impact of pathogenic mutations at these loci. Indeed, we found that high confidence risk genes for ASD (Sanders et al, 2015) are enriched two-fold for sex biased expression in the prenatal brain. The proportion of genes exhibiting sex-biased expression in the fetal brain was even higher for those implicated in developmental disorders by the recent Deciphering Developmental Disorders Study (2017), with the majority (13 /16) more highly expressed in males.

Analyses of our data in relation to recent single cell RNA sequencing findings (Fan et al. 2018; Zhong et al. 2018) suggest subtle differences in cellular composition between the male and female human brain during the studied period of gestation. Consistent with our finding that male-biased genes are enriched for involvement in the cell cycle, we find that these genes are also enriched for those that are more highly expressed in neural progenitor cells. In contrast, we observe a highly significant enrichment of female-biased genes among those that have been reported to be more highly expressed in Cajal-Retzius cells, as well as enrichments for genes that are prominently expressed in oligodendrocyte precursors and astrocytes. Cajal-Retzius cells are an early form of neuron that play a major role in neural migration through their secretion of the extracellular matrix protein reelin (Soriano & Del Rio. 2005), while oligodendrocyte precursor cells and astrocytes appear after the emergence of neurons in the mammalian brain (Qian et al. 2000; Zhong et al. 2018). It is possible that the cellular differences between the male and female prenatal brain suggested by these analyses have etiological relevance to ASD and other sex-biased neurodevelopmental disorders. For example, an extended period of neural progenitor proliferation might render males more susceptible to environmental and genetic insults, while earlier maturation of particular neuronal and glial cells in females might confer a protective effect. We provide our data as a searchable, online resource to aid in the investigation of these and other disorders of human brain development.

## METHODS

### RNA sequencing data

RNA sequencing data were previously generated using undissected brain tissue from 120 human fetuses aged 12-19 PCW (described in O’Brien et al. 2018). Briefly, RNA-Seq libraries were prepared from extracted total RNA using the TruSeq Stranded Total RNA RNA Library Prep kit (Illumina), following depletion of ribosomal RNA. Libraries were sequenced on Illumina HiSeq systems, generating at least 50 million read pairs (100 million reads) per sample. Fetal sex was determined by karyotyping, expression of genes on the Y-chromosome in males and heterozygosity for genetic X-chromosome markers in females. Of the 120 samples included in the analysis, 70 were thus determined to be male and 50 female. Male and female samples were well matched for known variables including age, RIN and read count (Supplementary Table S7).

### Gene expression analyses

Transcript abundance was quantified by pseudoalignment of sequencing reads to transcript sequences derived from the GRCh38 human genome reference sequence and Ensembl (version 81) reference annotation using kallisto (Bray et al. 2016). Reads were aggregated at the gene level using tximport (Soneson et al. 2015) and biomaRt (Durinck et al. 2005), with between-sample normalization and variance-stabilizing transformation carried out using DESeq2 (Love et al. 2014). Gene expression measures were quantile normalized and corrected for PCW, RIN, sequencing batch, the first three principal components derived from genome-wide DNA genotypes (O’Brien et al. 2018) and 10 hidden confounders estimated through use of probabilistic estimation of expression residuals (PEER; Stegle et al. 2012). Tests of differential expression between males and females were performed using DESeq2 (Love et al. 2014) using the wrapper scripts included in the SARtools package (Varet et al. 2016) in the R statistical computing environment. Genes and transcripts with low expression were filtered out using empirically determined thresholds (average counts per sample of 0.26 for the gene-level analysis and 0.23 for the transcript-level analysis) before controlling the False Discovery Rate (FDR) at 0.1. Outliers were detected by calculating Cook’s distance: genes where Cook’s distance was greater than 10 for any sample were discarded from the list of differentially expressed genes.

### Analyses of genes implicated in neurodevelopmental disorders

Enrichment of sex-biased genes (FDR < 0.1) among high confidence risk genes for ASD (Sanders et al. 2015) and developmental disorders (Deciphering Developmental Disorders Study, 2017) was tested using hypergeometric tests, as implemented in the clusterProfiler R package (Yu et al. 2010).

### Gene Ontology and cell enrichment analyses

Autosomal genes exhibiting sex-biased expression (FDR < 0.1) were subject to Gene Ontology (GO) enrichment analysis using all terms included in the comprehensive GOBPFAT category through DAVID Bioinformatics Resources 6.8 (Huang et al. 2009; https://david.ncifcrf.gov/). For each analysis, we used a background of all expressed autosomal genes (N = 30,331). To explore potential differences in cellular composition between the male and female prenatal brain, we tested for enrichment of male- and female-biased genes (FDR < 0.1) among genes that are reported by Zhong et al. (2018) to be differentially expressed between 6 major cell types in the prenatal human prefrontal cortex (neural progenitor cells, excitatory neurons, interneurons, astrocytes, oligodendrocyte progenitor cells and microglia) and among genes that are reported by Fan et al. (2018) to be differentially expressed between 8 major cell types of the human fetal cerebral cortex (excitatory neurons, inhibitory neurons, Cajal-Retzius cells, glial cells, microglia, endothelial cells, immune cells and blood cells). Enrichments were tested using hypergeometric tests, as implemented in the clusterProfiler R package (Yu et al. 2010), and resulting *P*-values were Bonferroni-corrected for 14 tests.

## DATA ACCESS

A searchable database of normalized expression data for all expressed Ensembl genes and transcripts can be accessed at: http://fgen.psycm.cf.ac.uk/FBSeq1. The raw RNA sequencing data are available through the European Genome-phenome Archive (https://www.ebi.ac.uk/ega/home) under study accession number EGAS00001003214. Scripts to produce analyses and figures are available at: https://github.com/hobrien/GENEX-FB1.

## ACKNOWLEDGEMENTS

This work was funded by a Medical Research Council (U.K.) project grant to NJB (MR/L010674/2). The human fetal material was provided by the Joint MRC/Wellcome Trust (grant #099175/Z/12/Z) Human Developmental Biology Resource (www.hdbr.org).

